# Systematic identification of pan-cancer single-gene expression biomarkers in drug high-throughput screens

**DOI:** 10.1101/2025.08.03.668348

**Authors:** Ginte Kutkaite, Göksu Avar, Diyuan Lu, Thomas J. O’Neill, Daniel Krappmann, Michael P. Menden

**Author notes:** Equal contribution (Ginte Kutkaite, Göksu Avar).

## Abstract

Precision oncology relies on molecular biomarkers to stratify patients into responders and non-responders to a given treatment. Although gene expression profiles have historically been explored for biomarker discovery, fewer studies investigated single-gene expression biomarkers. Additionally, many approaches are limited to cancer type-specific associations, which constrain statistical power. To address these limitations, we developed a regression-based framework that corrects for tissue-specific biases and enhances detection of pan-cancer single-gene expression biomarkers of drug sensitivity in cancer cell line high-throughput drug screens. Our method maintains predictive performance post-correction, and successfully recovers established biomarkers, such as *SLFN11* expression for DNA damaging agents. Notably, we identified *SPRY4* and *NES* expression as biomarkers of sensitivity for compounds targeting ERK/MAPK signaling (adjusted p-value=4.016×10⁻⁵ and 7.221×10⁻⁶, respectively). This approach offers a scalable strategy for biomarker discovery and holds potential for translation to more complex biological models and patient-derived datasets. Ultimately, pan-cancer single-gene expression biomarkers may improve patient stratification and clinical outcomes in precision oncology.

## INTRODUCTION

Precision oncology seeks to improve treatment outcomes by stratifying patients based on their molecular profiles to predict therapeutic response^1^. Despite advances in molecular profiling technologies, drug development remains high-risk, with clinical trial failure rates nearing 95%^2,3^ often due to the absence of reliable biomarkers for identifying responsive subgroups. This underscores the urgent need for novel biomarkers and innovative application strategies to accelerate drug development and improve clinical success^1^.

Biomarker discovery remains a major challenge in precision oncology. Large-scale efforts such as The Cancer Genome Atlas (TCGA)^4^ and the International Cancer Genome Consortium (ICGC)^5^ have mapped tumor molecular profiles, but largely lack linked treatment records and clinical outcomes. Real-world data (RWD) sources like Flatiron Health^6^ integrate molecular and clinical data from hospital cohorts but are limited by sparse coverage of investigational therapies, non-randomized treatment assignment, variable data quality, and restricted accessibility.

High-throughput drug screens in molecularly profiled cancer cell lines offer a scalable framework for biomarker discovery. Pioneering efforts such as the NCI-60 screen^7^ laid the foundation for larger-scale resources, including the Genomics of Drug Sensitivity in Cancer (GDSC)^8,9^ and the Cancer Cell Line Encyclopedia (CCLE)^10^, which profile drug responses in over 1,000 cancer cell lines spanning diverse tissue types. The integration of these data with multi-omics characterization supports the identification of pan-cancer biomarkers and mechanistic insights across genomic, transcriptomic, and epigenetic layers^9,11–13^.

A wide range of statistical and machine learning (ML) frameworks have been developed to model drug response in cancer cell lines, balancing predictive performance and interpretability. Biomarker discovery approaches span from univariate ANOVA models^9,11^ to multivariate regularized linear regression^12^. While more complex ML models, such as support vector machines, random forests, and deep neural networks, may offer higher predictive accuracy^14–16^, they often lack interpretability. To improve interpretability, post hoc model-agnostic methods such as Shapley values^17^ and LIME^18^ have been developed to quantify how individual features influence model predictions. Nonetheless, the biological limitations of cell line models underscore the need for rigorous biomarker validation in more clinically relevant systems.

Cancer is driven by genetic alterations, and accordingly, most drug response biomarkers are based on genetic alterations^19^. As one of the earliest and most extensively studied omic layers, genomics has yielded numerous mutation-based biomarkers across various cancer types^20,21^. While other omics, such as transcriptomics and proteomics, have also been widely used, their integration into systematic biomarker discovery efforts has been comparatively limited ^20,21^. Gene expression (GEX) signatures, also referred to as endotypes, are increasingly recognized for their association with drug response and are beginning to enter clinical practice^20,22^. However, single-GEX biomarkers are comparatively rare, partly due to their transient and context-dependent nature. A notable exception is *SLFN11*, whose upregulation sensitizes cancer cells to DNA-damaging agents and has been validated in preclinical models^10,23–25^.

GEX exhibits strong tissue-of-origin dependency, representing a major obstacle for the identification of pan-cancer biomarkers^26–28^. As a transient omic layer, it is governed by tissue-specific regulatory programs, leading to high consistency within tissues but poor comparability across them^29^. This tissue effect can confound associations with drug response, obscuring signals that generalize across cancer types. Consequently, many studies are restricted to single-tissue analyses, which limits statistical power and prevents leveraging cross-tissue or transfer learning opportunities.

Here, we aim to identify pan-cancer single-GEX biomarkers predictive of drug sensitivity. To reduce tissue-specific bias, we implemented two correction strategies: (1) z-score normalization and (2) residual adjustment. We then applied regularized linear regression to associate corrected single-GEX with drug response across cancer cell lines. Focusing on individual genes enables interpretable models, while the pan-cancer design increases sample size, statistical power, and transfer learning across cancer types. We hypothesize that this approach will recover known drug targets and uncover novel, clinically relevant biomarkers.

## RESULTS

For the discovery of pan-cancer single-gene expression (GEX) biomarkers, we first addressed tissue-of-origin effects in cancer cell lines. We analyzed GDSC data comprising 778 cell lines across 29 cancer types and drug response to 385 compounds targeting 24 pathways (**Fig. 1A**), with response quantified as area under the curve (AUC). Principal component analysis revealed strong tissue-specific expression patterns, particularly between solid and non-solid tumors (**Fig. 1B**; **Supplementary Fig. 1A**). To mitigate this, we applied z-score normalization and residual-based correction (**Methods**). Post-correction, tissue-specific clustering was no longer evident (**Fig. 1C**; **Supplementary Fig. 1B-D**), enabling a more robust and unbiased identification of pan-cancer expression biomarkers.

**Figure 1.**
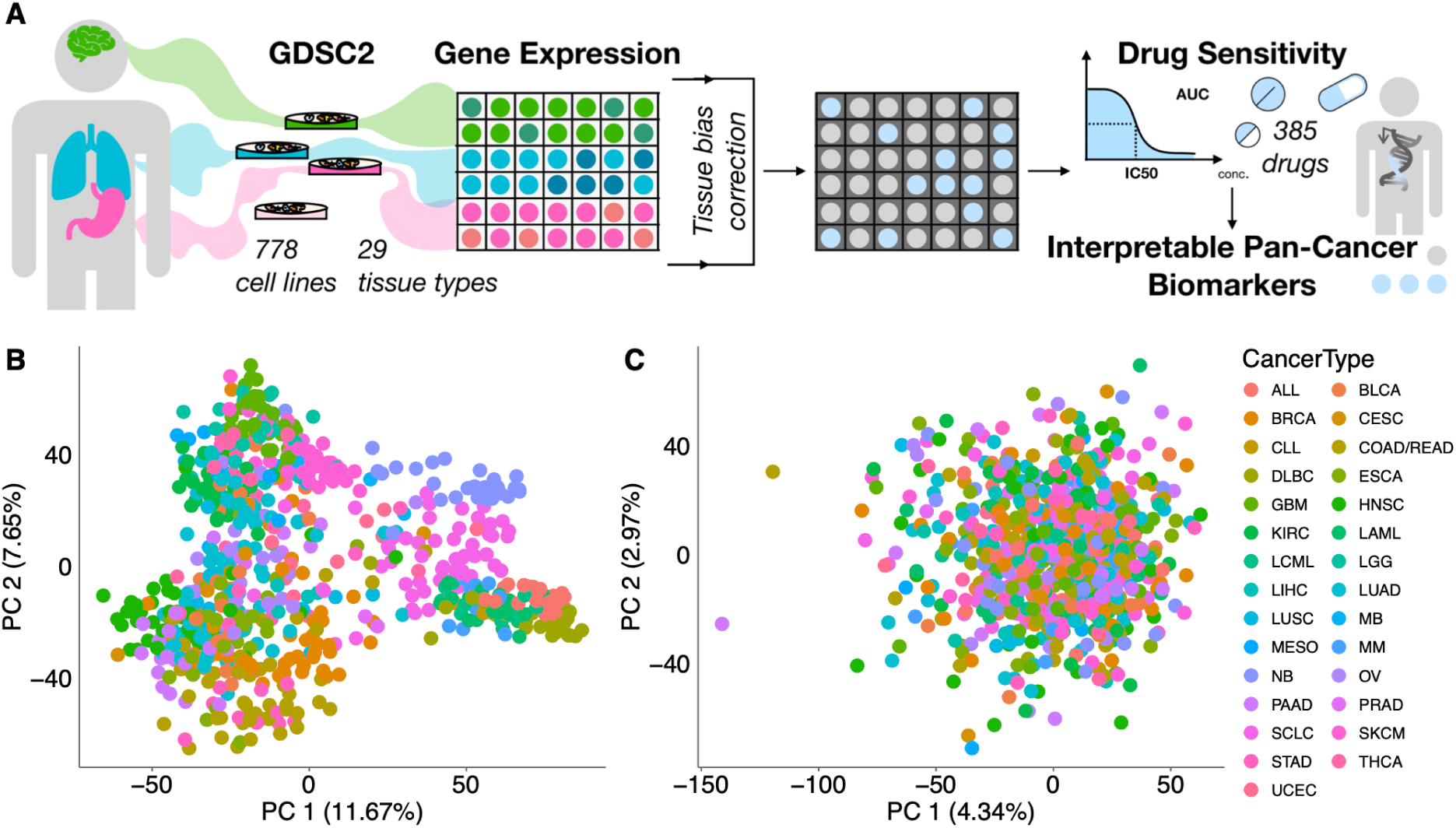
Tissue type dependencies in cancer cell line gene expression data. (**A**) Analysis workflow to identify gene expression biomarkers. (**B**) Principal Component Analysis (PCA) plot depicting gene expression data coloured by the cancer cell line tissue of origin. (**C**) PCA plot showing z-score corrected gene expression data.

### Disentangling tissue effects improves the accuracy of pan-cancer drug response predictions

To systematically predict drug response across the 385 compounds in a pan-cancer setting, we evaluated regularized linear regression models. While ridge regression underperformed relative to lasso and elastic net when using tissue labels as input, it outperformed both methods with GEX data (Wilcoxon signed-rank test adjusted p-value = 2.28×10⁻⁶⁴ vs. Lasso; 2.27×10⁻⁶⁴ vs. Elastic Net; estimate = 0.063; **Supplementary Fig. 2A-D**), and was therefore selected for all subsequent analyses (**Supplementary Table 1**). We trained 1,155 drug-specific ridge models using three input types: tissue labels (naïve baseline), uncorrected GEX (tissue-confounded), and tissue-corrected GEX (**Methods**; **Fig. 2A**).

**Figure 2.**
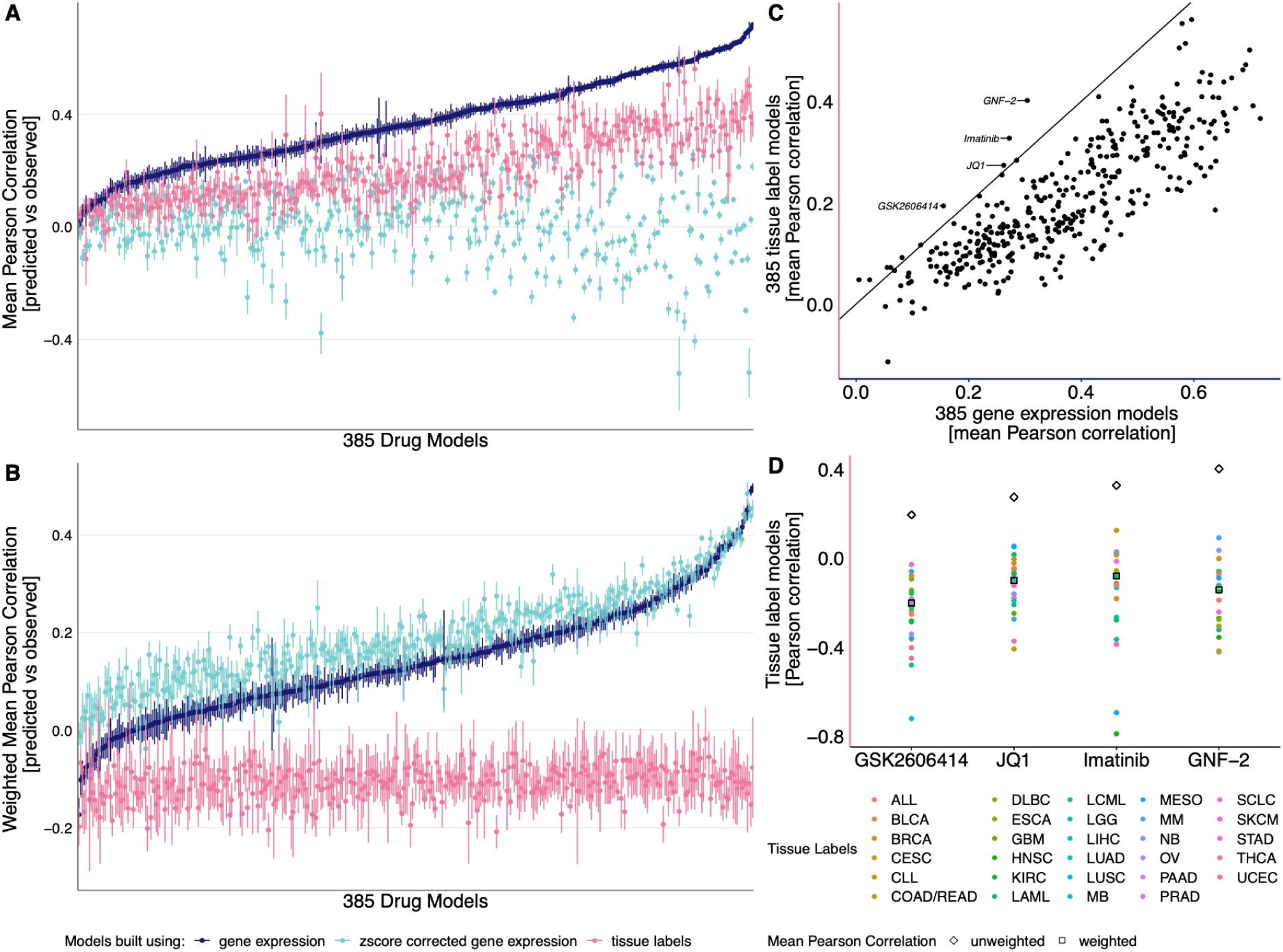
Performance of 385 drug models built using gene expression, z-score corrected gene expression and tissue labels. (**A**) Unweighted and (**B**) weighted Pearson correlation of 385 drug response models either leveraging gene expression, z-score corrected gene expression or tissue labels. (**C**) Mean unweighted Pearson correlation of drug models using tissue labels and gene expression. (**D**) Pearson correlation within individual tissue types as well as mean unweighted and weighted Pearson correlation.

A key challenge is Simpson’s paradox, arising from strong tissue-specific drug responses, for instance, cell lines derived from non-solid tumors often require lower drug concentrations than solid tumors, inflating prediction-observation correlations (**Supplementary Fig. 3**). Without bias correction, tissue labels and uncorrected GEX appear highly predictive (**Fig. 2A; Supplementary Fig. 2E**). Consequently, tissue-corrected GEX models seem to underperform (p-value<2.2×10⁻¹⁶; pseudo-median=-0.358 vs. GEX, pseudo-median=-0.197 vs. tissue; **Fig. 2A**), although this reflects non-translatable tissue effects with limited clinical utility.

To account for confounding due to tissue-specific drug responses and imbalanced tissue representation, we evaluated model performance using tissue-weighted Pearson correlation (**Methods; Fig. 2B**). This adjustment effectively removed the predictive advantage of models relying solely on tissue labels, which subsequently performed at random or at overfitted levels (**Fig. 2B; Supplementary Fig. 2F**). Notably, uncorrected GEX retained most its predictive power, but was significantly outperformed by tissue-corrected GEX models (p-value<2.2×10⁻¹⁶, pseudo-median=0.048), highlighting their value for pan-cancer drug response prediction (**Fig. 2B**).

Gene expression (GEX) encodes both tissue-of-origin and additional mechanistic information^30^, evidenced by GEX-based models generally outperforming tissue-based models in predicting drug response (**Fig. 2A**). We identified 10 exceptions where tissue labels yielded better performance (**Fig. 2C**), with four models exceeding a correlation of 0.15. These cases suggest that tissue origin may act as a proxy biomarker, and GEX models may overfit without adding mechanistic insight. Notably, certain cancer types are defined by genetic alterations; for example, imatinib and GNF-2, which target ABL, performed best in BCR-ABL-positive tissues characteristic of chronic myeloid leukemia (CML)^31^. However, this association is highly tissue-dependent and entirely lost within cancer type context (**Fig. 2D; Supplementary Fig. 2G-J**). These findings underscore the importance of not relying solely on the tissue context when refining predictive modelling and guiding biomarker discovery.

### Feature extraction allows pan-cancer gene expression signature discovery

Patient stratification in clinical settings necessitates the extraction of interpretable biomarkers of drug response. Here, we focused on predictive models derived from cancer cell line data, which serve as a preclinical framework for identifying such biomarkers. Ensuring robust model performance is essential to guarantee that selected features reflect meaningful biological signals. To systematically identify such models, we constructed null models and applied standard deviation-based thresholding, yielding 266 informative drug models spanning 23 distinct pathways (**Methods; Supplementary Fig. 4**).

Recurrent biomarkers shared across drugs targeting the same pathway may reflect underlying mechanistic associations. We investigated this by analyzing feature overlap across the 266 informative models at the pathway level (**Fig. 3A; Methods**). Models targeting EGFR signaling exhibited the fewest unique top-ranked features, suggesting strong recurrence of specific genes among the top 10 features across these drugs. To quantify this, we identified features that appeared in the top 10 for at least 25% of drugs targeting the same pathway (**Fig. 3B**), highlighting candidates with potential pathway-level relevance.

**Figure 3.**
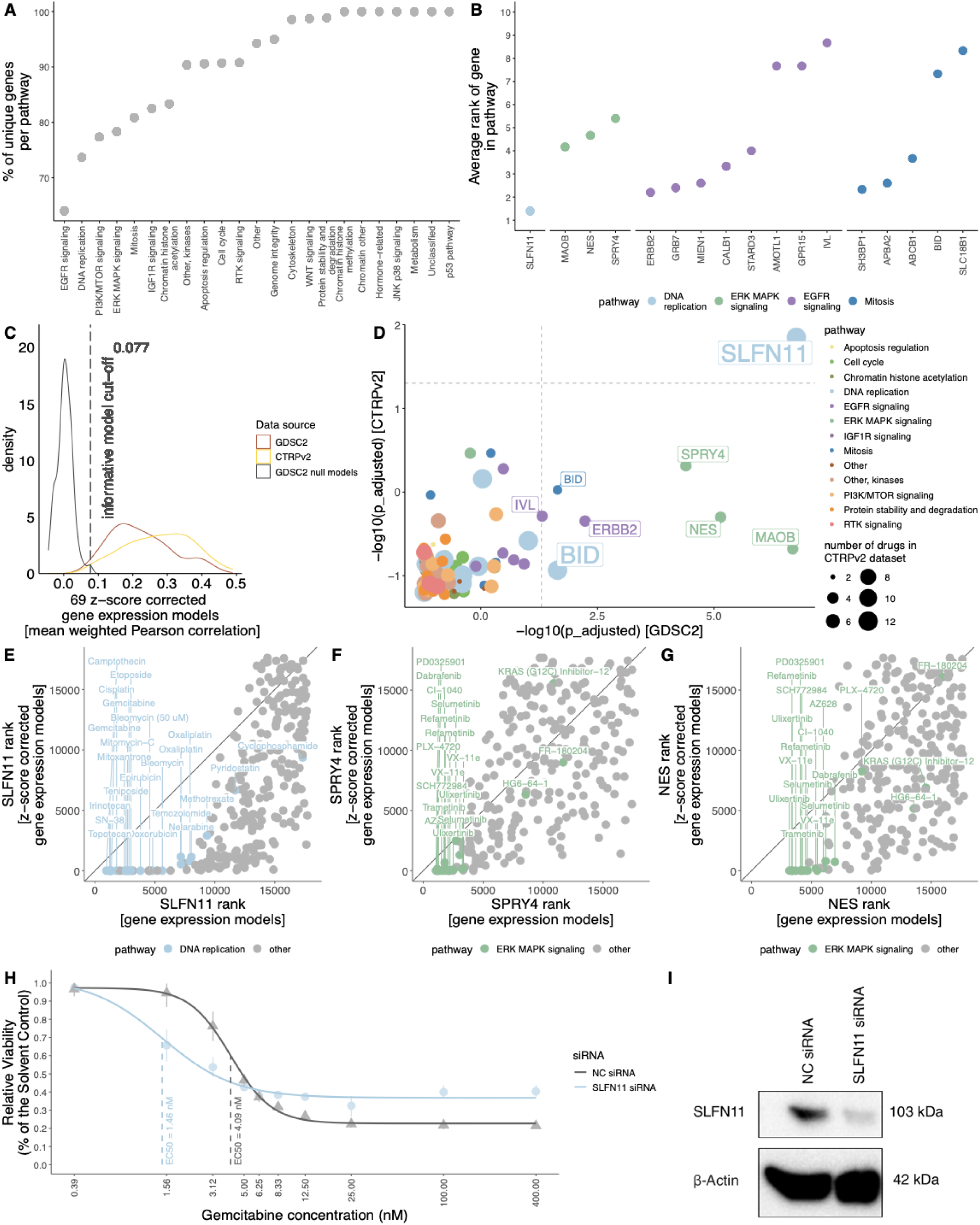
Robust pan-cancer biomarker selection. (**A**) The percentage of unique genes per pathway; (**B**) the average rank of genes for drugs in a specific pathway; (**C**) distribution of performance of 69 drug models built using the GDSC and the CTRP datasets; (**D**) adjusted p-values from overrepresentation tests with the GDSC and the CTRP datasets (points with negative enrichment score coloured by pathway); the rank of (**E**) *SLFN11*, (**F**) *SPRY4,* and (**G**) *NES* in models built with gene expression and z-score corrected gene expression inputs. (**H**) Dose-response curves of A375 melanoma cells transfected with *SLFN11* and negative control (NC) siRNAs and treated with gemcitabine for 72 hours. Dots and whiskers represent the mean viabilities with 95% confidence interval. (**I**) Western blot showing the decrease in SLFN11 protein levels 72 hours after transfection with siRNA.

Independent validation is essential to assess the robustness and translational potential of identified biomarkers. To this end, we used the Cancer Therapeutics Response Portal (CTRP) dataset, which provided matching drug response and GEX data for 69 of the 266 informative drugs identified in GDSC (**Supplementary Fig. 5; Supplementary Table 2**). Model performance showed significant concordance between the two datasets, particularly for tissue-corrected GEX models (Wilcoxon signed-rank test: GDSC vs. CTRP, p-value=9.688×10⁻⁵, pseudo-median=-0.048; GDSC null vs. CTRP, p-value=5.331×10⁻¹³, pseudo-median=-0.275; Fig. 3C), supporting the reproducibility of our modeling framework across an independent dataset.

To systematically assess pathway-level biomarker recurrence, we performed hypergeometric enrichment analysis to identify genes consistently ranked among the top features in drugs targeting the same pathway (**Methods**). This analysis confirmed *SLFN11*, a well-established and widely validated biomarker for sensitivity to DNA-damaging agents, as a recurrent feature in DNA replication-targeting drugs across both datasets (Fig. 3D**; Supplementary Fig. 6B-C**), serving as a positive control that supports the validity of our approach. Although the smaller sample size in CTRP (**Supplementary Fig. 5C-D**) limited replication of all top associations from GDSC, it nevertheless enabled the recovery of key biomarkers with consistent trends (Fig. 3D), yielding a set of candidates for further investigation.

In support of this, *SLFN11* was selected in 68% of DNA replication-targeting models (**Supplementary Fig. 6A**), with an average rank of 1.4 using z-score-corrected GEX (Fig. 3B). It was significantly overrepresented in both datasets following tissue bias correction (ES GDSC=-0.932, adjusted p-value=1.718×10⁻⁷; ES CTRPv2=-0.892, adjusted p-value=0.014; Fig. 3D**-E****; Supplementary Fig. 6B-C**), further supporting the robustness and reproducibility of our findings.

We next examined recurrent gene expression biomarkers associated with other drug-targeted pathways beyond DNA replication. Within the ERK MAPK signaling pathway, *SPRY4* and *NES* emerged as notable candidates. *SPRY4* and *NES* were selected in 28% and 33% of models, respectively, with average ranks of 5.4 and 4.7 (**Supplementary Fig. 6A**; Fig. 3B). Both genes were significantly overrepresented in the GDSC dataset after tissue correction (*SPRY4*: ES=-0.911, adjusted p-value=4.016×10⁻⁵; *NES*: ES=-0.908, adjusted p-value=7.221×10⁻⁶; Fig. 3D, F**-G**; **Supplementary Fig. 6B**), suggesting their potential as pathway-specific biomarkers.

We also recovered *ERBB2* in models targeting EGFR signaling, consistent with its well-established role as a biomarker supported by multiple studies^32–35^ (Fig. 3B, D**; Supplementary Fig. 6A-C, E**). In addition, other promising candidates emerged, including *MAOB* for ERK MAPK signaling, *IVL* for EGFR signaling, and *BID* for mitosis and DNA replication-targeting drugs (Fig. 3B, D**; Supplementary Fig. 6**). These findings highlight a broader spectrum of gene expression biomarkers that may warrant further functional validation and investigation.

Finally, we sought experimental validation of *SLFN11* as a gold standard biomarker to confirm the reliability of our computational framework. Given its well-established role in sensitizing cells to DNA-damaging agents, as well as consistent associations across GDSC and CTRP datasets, *SLFN11* was prioritized for *in vitro* validation over newly identified candidates. In *SLFN11*-knockdown A375 melanoma cells treated with gemcitabine, a DNA replication-targeting drug, we observed reduced drug efficacy upon *SLFN11* downregulation (EC50 NC=4.09nM vs EC50 SLFN11=1.46nM; Fig. 3H**-I****; Supplementary Fig. 6I**). This result not only aligns with known biology but also demonstrates that our framework can identify biomarkers with strong mechanistic and translational relevance.

## DISCUSSION

Genomic profiling within individual cancer types has driven early success in precision oncology by enabling targeted therapies against recurrent oncogenic mutations. However, progress has slowed due to tumor heterogeneity, limited cohort sizes, and the rarity of actionable mutations, all of which constrain predictive modeling and clinical translation. In contrast, gene expression (GEX) profiling and pan-cancer analyses remain underutilized^20–22,36^, despite their potential to capture functional tumor states and offer increased statistical power. Harnessing these complementary data layers presents a key opportunity to accelerate progress in precision oncology.

Cancer cell lines offer a scalable model for drug response studies, enabling experiments not feasible in patient-derived samples. Large-scale screens such as NCI-60, GDSC, and CTRP have validated known biomarkers and identified novel ones using statistical and machine learning methods^7,9–13^. Tissue-specific models often miss biomarkers in rare cancer types due to limited sample representation^9,11,12^. Pan-cancer approaches improve predictive performance but may obscure biological mechanisms, as they group distinct diseases that, despite shared hallmarks, differ in molecular pathogenesis^14–16,37,38^.

This study advances current computational approaches by systematically leveraging gene expression data in cancer cell lines to identify robust pan-cancer single-gene biomarkers. Our framework enables deeper insights into drug mechanisms of action and enhances the clinical translatability of biomarker discoveries. Notably, models incorporating tissue type-corrected gene expression retain strong predictive performance while yielding biologically interpretable biomarkers (Fig. 2B; Fig. 3D).

Effective correction of tissue-specific bias is essential for identifying meaningful pan-cancer biomarkers from gene expression data. We evaluated two correction strategies: residual-based and z-score normalization. Both approaches reduced tissue-driven variation, but z-score normalization provided a more stringent correction (Fig. 1C**; Supplementary Fig. 1B-D**). In contrast, residual-based correction retained subtle tissue-specific signals, as reflected in residual clustering when comparing solid vs non-solid tumor types (**Supplementary Fig. 1D**). Based on these observations, we used models trained on z-score normalized expression data for downstream biomarker interpretation.

Robust biomarker discovery should recover established associations and reveal biologically plausible candidates across diverse drug classes. In our analysis, the strongest biomarker signals were observed for compounds targeting DNA replication, ERK MAPK signaling, EGFR signaling, and mitosis (Fig. 3B). As expected, we recapitulated well-characterized biomarkers, including *ERBB2* for EGFR-targeting agents^32–35^ and *SLFN11* for DNA replication inhibitors^10,23–25,39–41^. This aligns with our validation experiment that siRNA-mediated *SLFN11* downregulation in A375 melanoma cells leads to reduced gemcitabine efficacy (Fig. 3H**-I**), further supporting the role of *SLFN11* as a key marker for sensitivity to DNA replication-targeting therapies.

ERK/MAPK pathway activity emerged as a key determinant of drug response in our analysis (Fig. 3B). Expression of *SPRY4* and *NES* correlated with sensitivity to ERK/MAPK pathway inhibitors. *SPRY4* encodes a known negative regulator of MAPK signaling via inhibition of GTP-bound RAS formation^42–44^. Although *SPRY4* has not previously been reported as a single-gene biomarker, it contributes to the MAPK Pathway Activity Score (MPAS), a transcriptional signature predictive of MEK1/2 inhibitor response in multiple cancer types^45^. Our findings support *SPRY4* expression as a potential surrogate marker of ERK/MAPK pathway activity and drug sensitivity.

*NES* serves as an additional gene expression biomarker of sensitivity to ERK/MAPK pathway inhibitors. NES is an intermediate filament protein that facilitates mitotic progression by disassembling phosphorylated vimentin^46–48^. It is recognized as a cancer stem cell marker^49^ and promotes tumor proliferation and invasion via mitochondrial remodeling^50^. Supporting our findings, *NES* expression in melanoma has been linked to increased sensitivity to BRAF and MEK inhibitors, including dabrafenib and trametinib^51^.

In conclusion, our pan-cancer gene expression analysis of cancer cell lines identified both known and novel drug sensitivity biomarkers, including *SPRY4* and *NES* for ERK/MAPK pathway inhibitors. This approach offers a scalable framework for generating biomarker hypotheses across diverse drug classes. This strategy can be readily extended to other preclinical models, including patient-derived organoids and xenografts, to better capture tumor heterogeneity and improve clinical translatability. Ultimately, such integrative analyses may inform the development of systematic, biomarker-guided strategies for tailored treatment selection in oncology.

## MATERIALS and METHODS

### Cancer Cell Line Characterization

Robust Multichip Average (RMA) normalized gene expression data as well as annotations such as MSI status, growth properties and culture media information for 1,001 cell lines can be downloaded from GDSC portal (https://www.cancerrxgene.org/downloads).

From 1,001 cell lines, 223 were filtered out as they did not have full information, namely, gene expression, consistent tissue labels or drug response data. Therefore, models were built on 778 cancer cell lines.

### Drug response data

The drug response data can be downloaded from the Genomics of Drug Sensitivity in Cancer (GDSC) portal data release 8.0 (https://www.cancerrxgene.org/downloads). Where available, we have used GDSC2 data. Drug response was quantified by area under the drug response curve (AUC).

Out of 400 tested drugs 15 were filtered out where a given drug was tested in fewer than 10% of all cell lines(*n* = 778) or tested in only one specific cancer type, leaving 385 drugs to be used in building pan-cancer models.

Cancer Therapeutics Response Portal (CTRP) validation data

For validation of our pan-cancer biomarkers, we have used drug response (AUCs) and gene expression data downloaded from CTRPv2 via National Cancer Institute portal (https://ctd2-data.nci.nih.gov/Public/Broad/CTRPv2.1_2016_pub_NatChemBiol_12_109/).

Out of 481 drugs in the dataset, 69 were overlapping with our informative drugs (**Methods**; **Supplementary Fig. 5C**) and had corresponding gene expression information. Models for these drugs were built using gene expression and z-score corrected gene expression matrices (**Methods**) from 822 cell lines across 23 cancer lineages (**Supplementary Fig. 5A-B**).

### Drug response predictions

To predict drug response and ultimately retrieve pan-cancer biomarkers, we have employed linear regression models, namely ridge^52^, lasso^53^ and elasticNet^54^, from glmnet R package. The fundamental difference between these models is the values of the tuning parameter alpha (0 < α < 1). with Ridge defined by α = 0 and Lasso defined by α = 1. For elasticNet we have tested alphas of 0.2, 0.4, 0.5, 0.6 and 0.8 with tissue label as well as gene expression matrices as input (**Supplementary Fig. 2B, D**). No significant difference in model performances was noted between different alpha parameters.

For all models, 10-fold cross-validation is applied and repeated 10 times. The weighted and unweighted Pearson correlations were used to evaluate the model performance. The weighted Pearson correlation (*pw*) was calculated as follows,

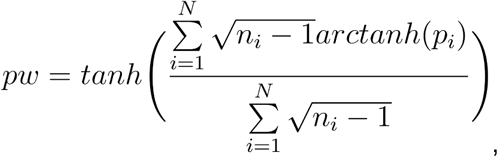

where *i* is an individual cancer type, *N* = 29 is the number of tested cancer types and *p_i_* is an unweighted Pearson correlation. For a given tissue type and drug combination, at least 3 cell lines had to be treated (*n_i_* ≥ 3).

### Z-Score-based tissue-correction

To account for the difference between tissues, gene expression data is normalized by subtracting the tissue-specific mean and divided by the tissue-specific standard deviation. The z-score is calculated as shown below

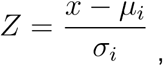

where *μ_i_* and *σ_i_* are the mean and the standard deviation of tissue type *i* (*i* = 29) across all cell lines, respectively. *x* is a RMA normalized gene expression count. Unless specified otherwise, all results reported to tissue-corrected GEX are based on the z-score correction.

### Residual-based tissue-correction

Additionally to the z-score-based correction, we built generalized linear models to predict gene expression profiles from the tissue type labels alone, followed by residual extraction. The procedure was repeated ten times and an averaged residual matrix was subsequently used to predict drug dose response in pan-cancer. Yielding similar results to z-score based correction, these results were reported in the supplements (**Supplementary Fig. 1C-D**, **4C**, and **Supplementary Table 2**).

### Null models and thresholding

In order to select informative models, a suitable performance threshold is needed. To select this threshold, we have built null models with shuffled drug dose-response data. For each drug model (*n* = 385), we have generated ten null-matrices, which serve as the drug response baseline for predictions.

A distribution of null models with mean-weighted Pearson correlation values was built. The performance threshold is defined as the mean plus three standard errors of the null model distribution. This was used to select informative models for each input matrix, namely, tissue labels, gene expression, residual– and z-score-corrected gene expression (**Supplementary Fig. 4A-D**). We have annotated 266 models as overall informative (**Supplementary Fig. 4E-F**).

### Feature selection and processing

To better understand the biological implication, we have further investigated the features of those selected informative models (*n* = 266). Consider the total number of genes is *G*. For drug *d*, 10 independent runs were performed and the weights for each gene *g* in all 10 runs were collected and averaged, denoted by 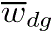. Then, the averaged weight 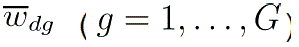 was sorted by their absolute values in a descending manner across all genes. This gave rise to the average rank of gene *g* for drug *d*, denoted by 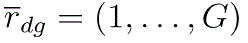, which is the index of 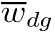 in the sorted weight vector. This was repeated for all the informative drug models (*n* = 266).

To summarize feature information on a pathway level, we focused on those drug models that target the same pathway. Given a total number of *d_p_* drug models targeting pathway *p*, we only considered the top 10 ranked genes in each drug model, i.e., 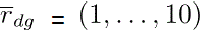, resulting in a total number of 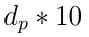 genes (one gene might appear multiple times). Then, the percentage of the unique genes for pathway *p* was computed by

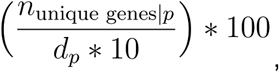

where 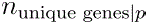 is the number of unique genes for pathway *p*.

### Hypergeometric enrichment analysis

We ran an enrichment analysis to systematically identify which features are overrepresented with high ranks in certain drug pathways. To this end, we leveraged the fgsea function from fgsea R package with a vector of drugs ranked from lowest to highest weight for each feature of interest (*n* = 17, selected from Fig. 3B). Here, only pathways targeted by at least two drugs were considered. The enrichment scores (ES), p-values as well as the Bonferroni p-adjusted values were estimated for each feature and pathway combination.

### Cell culture

A375 melanoma cells (source: ATCC) were cultured in Gibco Dulbecco’s Modified Eagle Medium supplemented with 10% Fetal Bovine Serum and 1% Penicillin-Streptomycin (10000 U/mL) in a humidified incubator (37°C, 5% CO2).

### siRNA mediated knockdown

10.000 A375 melanoma cells per well were reverse transfected in an opaque, white, flat-bottom plate, using *SLFN11* Silencer Select Pre-designed siRNA (Ambion: 4392420) and Silencer Negative Control siRNA #1 (Ambion: AM4611) with Lipofectamine RNAiMAX transfection reagent (Invitrogen: 13778075) and Gibco Opti-MEM reduced serum medium, following the manufacturer’s protocol for 1.5 pmol siRNA per well.

### Gel Electrophoresis and Western blotting

Cells were lysed using co-immunoprecipitation buffer (150mM NaCl, 25mM HEPES, 0.2% NP40, 1mM Glycerol) supplemented with cOmplete Protease Inhibitor Cocktail (Roche: 11836145001) and the protein concentrations were analyzed using Quick-Start Bradford 1X Dye Reagent (Bio-Rad: 5000205). The proteins were detected using anti-beta-Actin Antibody C4 (Santa Cruz: sc-47778) and anti-SLFN11 antibody (Abcam: ab121731).

### Drug treatment and dose response analysis

After the transfected cells were incubated overnight, they were treated with Gemcitabine (SelleckChem: S1714) dissolved in DMSO (0.5% v/v DMSO concentration per well). 72 hours after the treatment, cell viability was measured using CellTiter-Glo 2.0 Cell Viability Assay (Promega: G924A). Relative viability as a percentage of the negative control was calculated with intensities from blank (I_B_: medium only), negative control (I_NC_: DMSO treatment) and Gemcitabine treatment (I_G_) wells as:

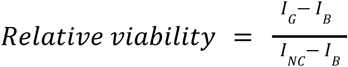

Dose-responses were analyzed using the four-parameter log-logistic (LL.4) model in the R package ‘drc’ ^55^.

## ACKNOWLEDGEMENTS

We are grateful for the valuable discussions with colleagues at Helmholtz Munich and the support from our funding agencies.

## AUTHOR CONTRIBUTIONS

Conceptualization, M.P.M.; Data curation, G.K.; Supervision, M.P.M. and D.K.; Formal analysis, G.K.; Methodology, G.K., M.P.M., T.J.O., and D.K.; Designing and performing experiments, G.A., T.J.O., and D.K.; Visualization, G.K.; Writing-original draft, G.K.; Writing-review and editing, G.K., D.L., G.A., T.J.O., D.K. and M.P.M. All authors have read and agreed to the published version of the manuscript.

## SUPPLEMENTARY FIGURES

**Supplementary Figure 1.**
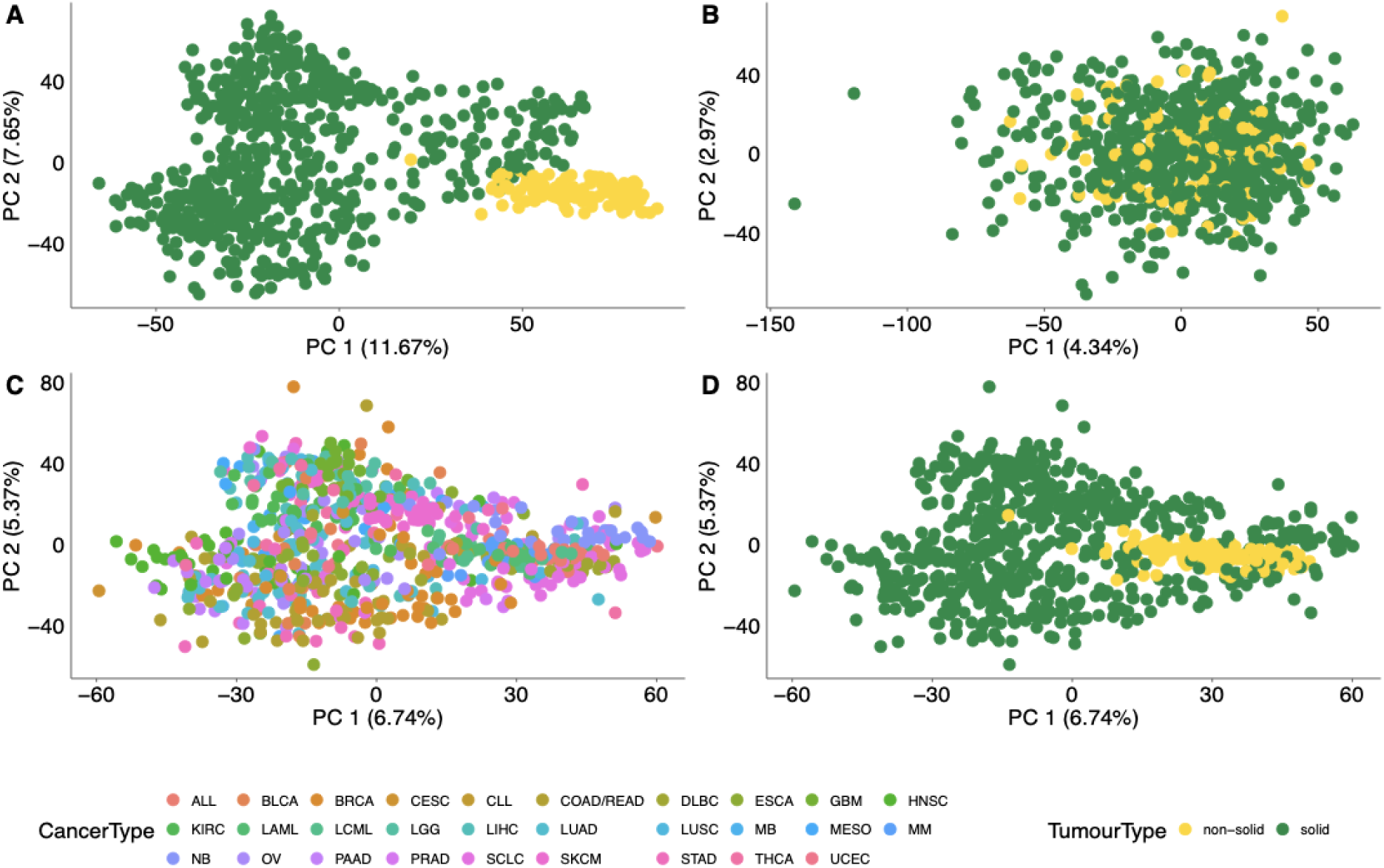
Principal Component Analysis plots and heatmap depicting gene expression data. (**A**) Gene expression data; (**B**) z-score corrected gene expression data; (**C**) residual corrected gene expression data (colored by cancer types); (**D**) residual corrected gene expression data (colored by cancer tumor types).

**Supplementary Figure 2.**
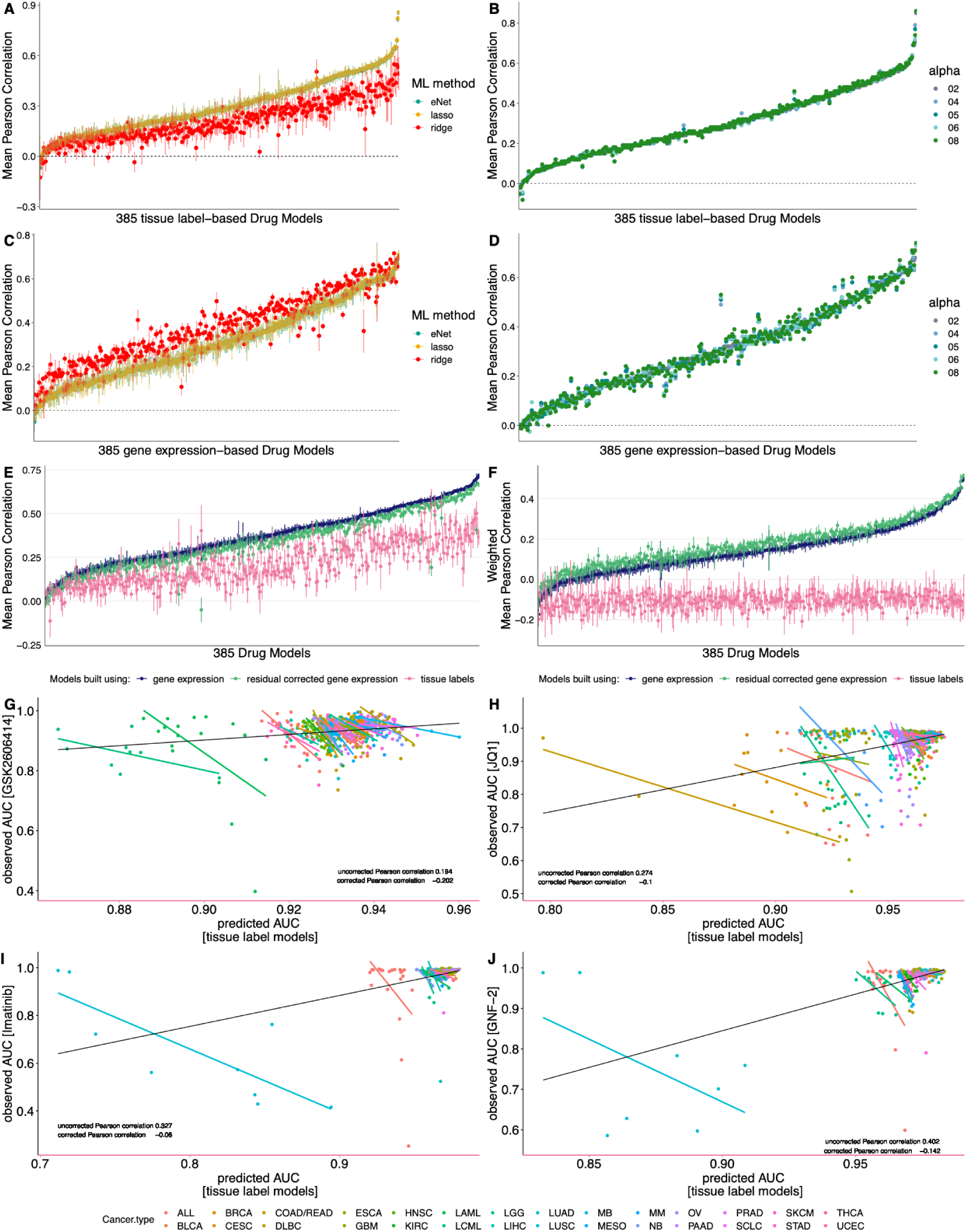
Method selection and model performance. (**A**) Performance of models built with ridge, lasso and (**B**) elasticNet regressions using tissue label data; (**C**) performance of models built with ridge, lasso and (**D**) elasticNet regressions using gene expression data; (**E**) Unweighted and (**F**) weighted Pearson correlation of models built with tissue labels, gene expression and residual corrected gene expression; observed and predicted AUC values with tissue labels models for (**G**) GSK2606414, (**H**) JQ1, (**I**) imatinib and (**J**) GNF-2.

**Supplementary Figure 3.**
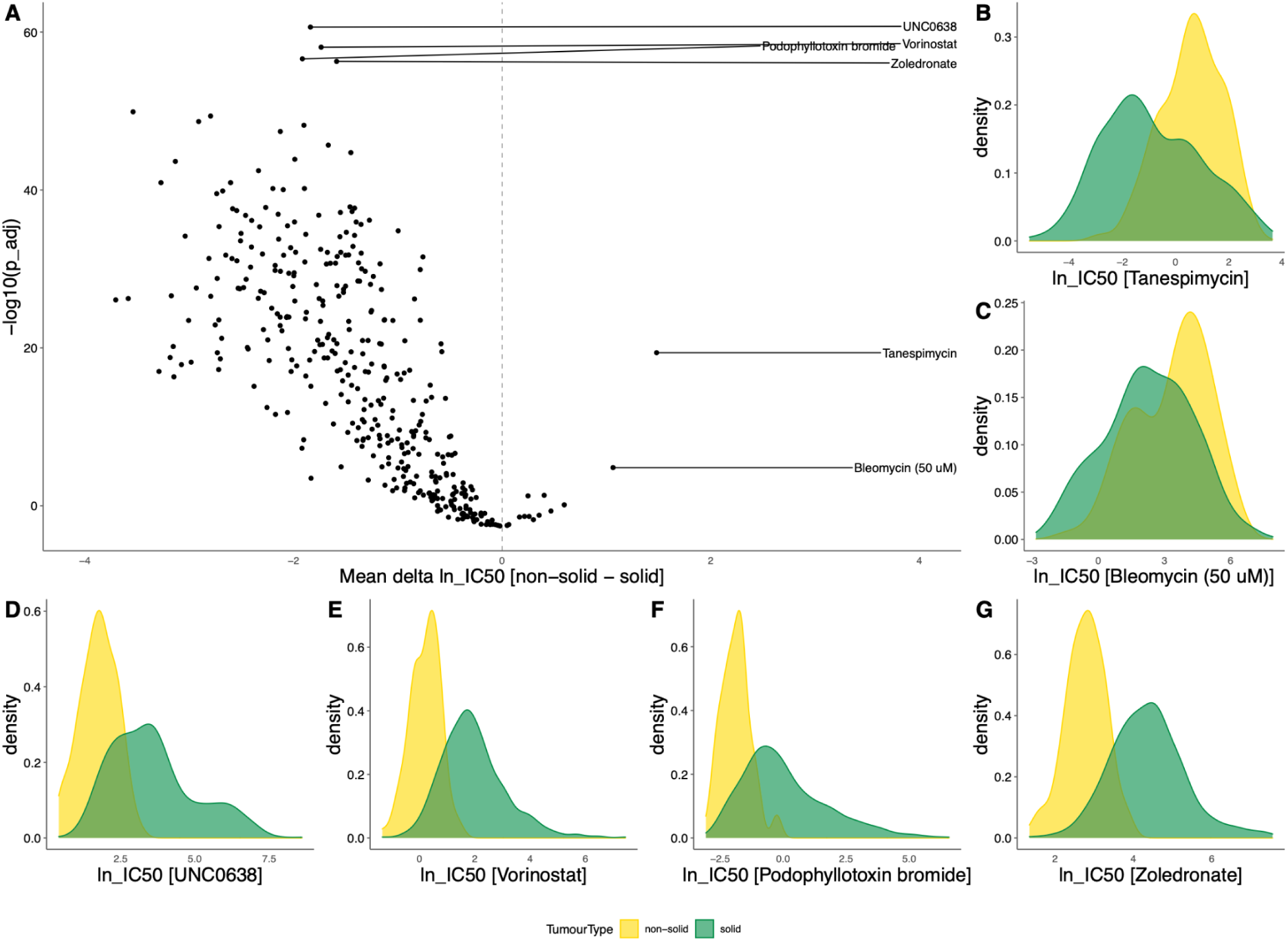
Difference in drug response IC50s between non-solid and solid tumor types. (**A**) Mean difference between drug (n=385) IC50s; density plots of IC50 of (**B**) tanespimycin, (**C**) bleomycin, (**D**) UNC0638, (**E**) vorinostat, (**F**) podophyllotoxin bromide, and (**G**) zoledronate.

**Supplementary Figure 4.**
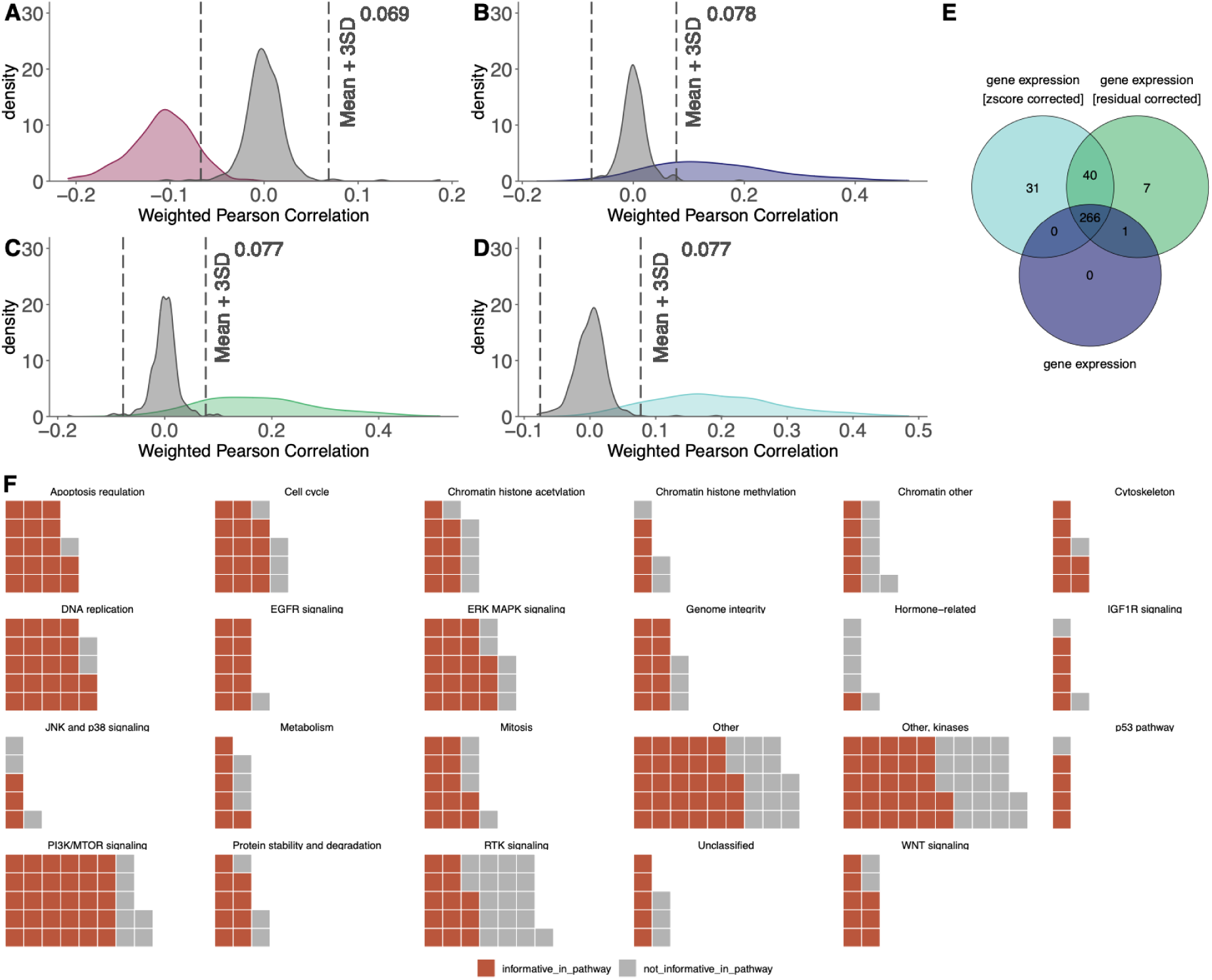
Model selection. Distribution of weighted Pearson correlation of models built with (**A**) tissue labels alone (**B**) gene expression (**C**) residual corrected gene expression and (**D**) z-score corrected gene expression as well as respective null models (grey). (**E**) Overlap of informative models built using different modalities. (**F**) Overview of drug models which were classified as informative (n=266) (not-informative in grey) stratified by pathway.

**Supplementary Figure 5.**
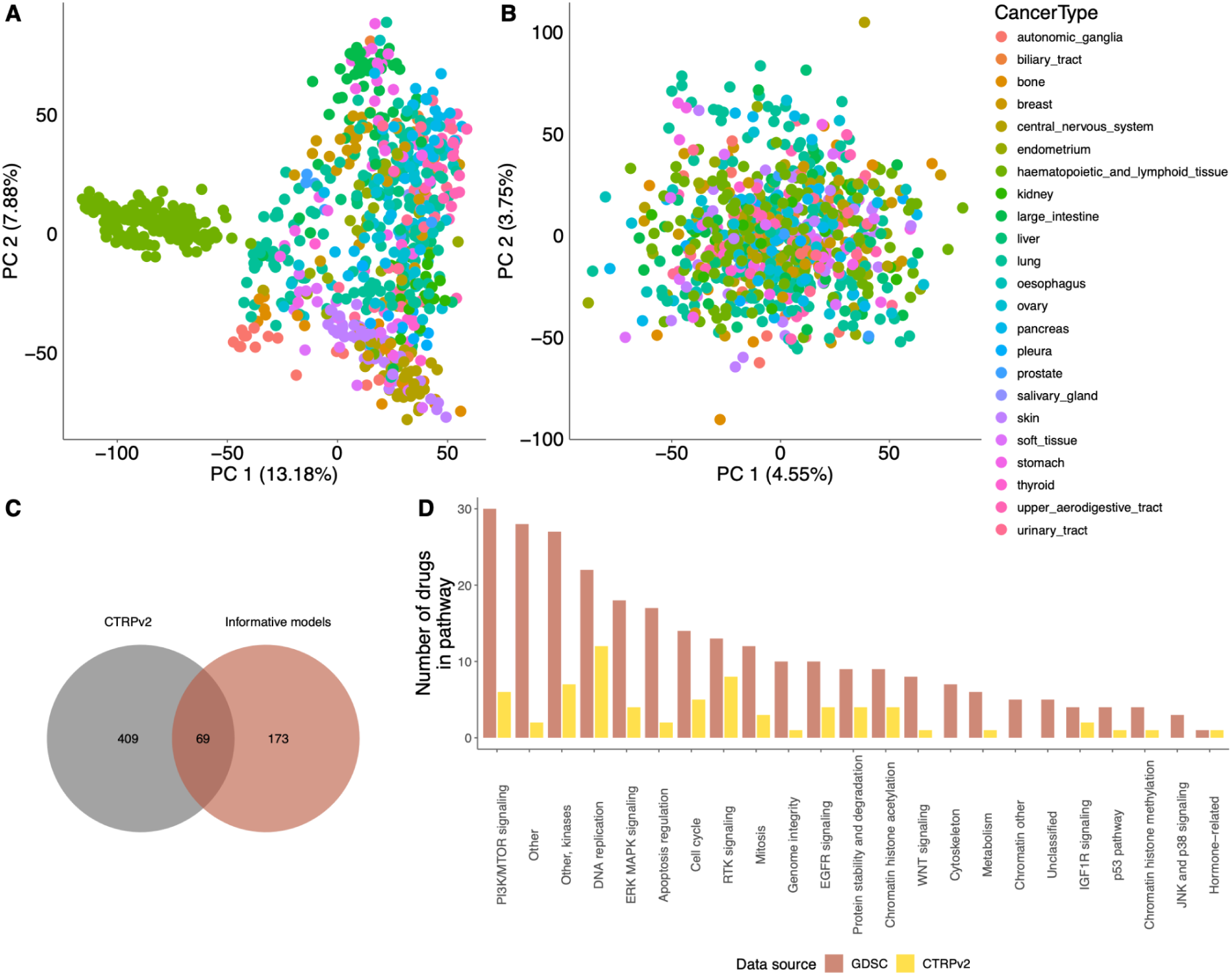
CTRP validation dataset. (**A**) Principal Component Analysis (PCA) plot depicting gene expression data coloured by the tissue origin of the cancer cell lines; (**B**) PCA plot depicting z-score corrected gene expression data; (**C**) an overlap of informative drug models with drugs screened in CTRP dataset; (**D**) number of drugs per pathway stratified by data source, GDSC (n=266) and CTRP (n=69).

**Supplementary Figure 6.**
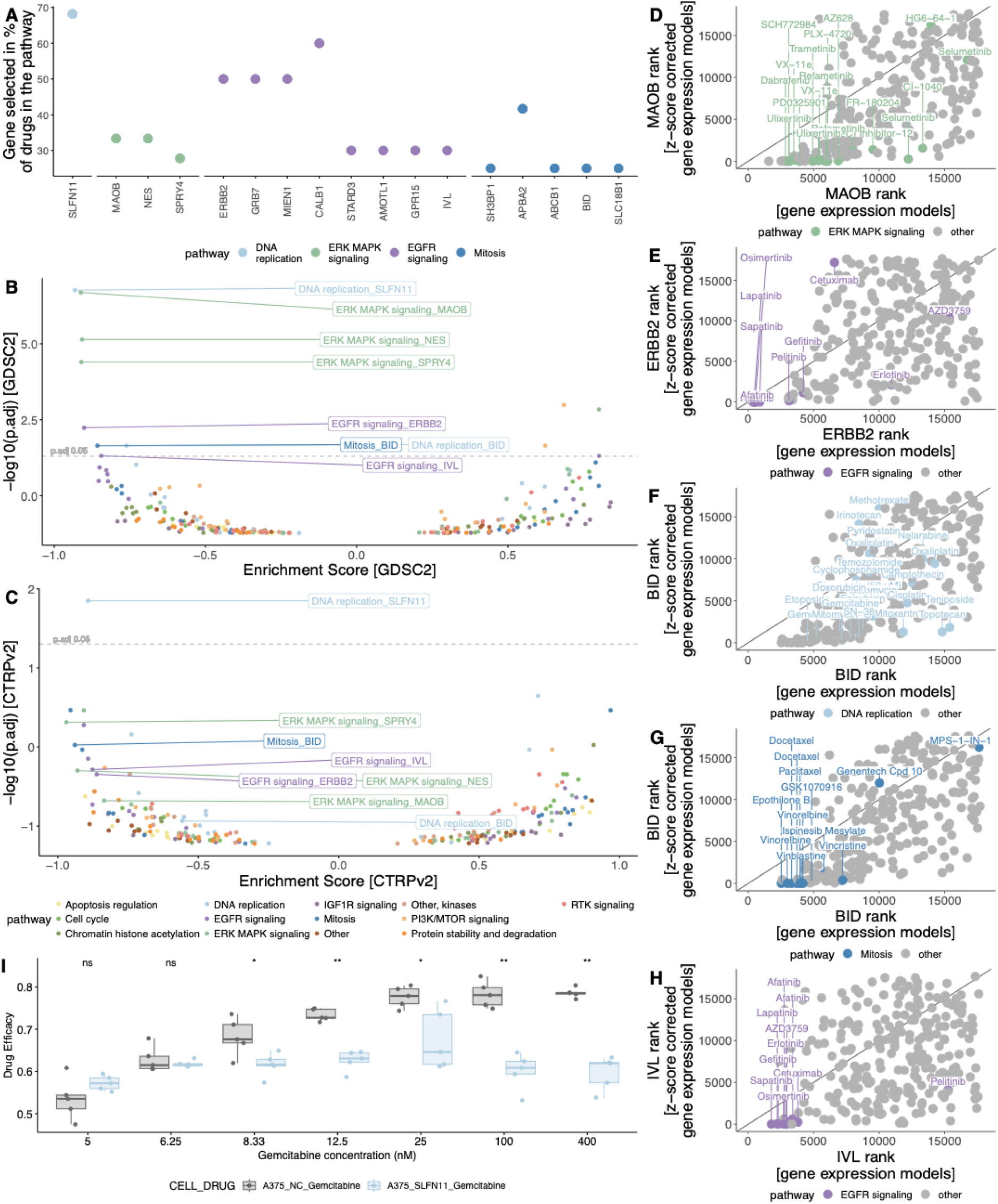
Robust pan-cancer biomarkers. (**A**) Percentage of drugs within pathway where specific gene is ranked in the first 10 positions; volcano plot of genes (n=17) enriched in (**B**) GDSC and (**C**) CTRP drug pathways; rank of (**D**) *MAOB*, (**E**) *ERBB2*, (**F**) *BID* with DNA replication targeting drugs, (**G**) *BID* with mitosis targeting drugs, and (**H**) *IVL* in models built with gene expression and z-score corrected gene expression inputs. (**I**) Efficacy of Gemcitabine on *SLFN11* knockdown (blue) and negative control (grey) A375 melanoma cells. Wilcoxon test, ns: p > 0.05,*: p <= 0.05,**: p <= 0.01.

## DATA AVAILABILITY

1. Supplementary Table 1. Ridge, lasso and elasticNet gene expression model performance.
2. Supplementary Table 2. Informative model (n=266) built on gene expression, z-score-, and residual-corrected gene expression performances.

## CODE AVAILABILITY

The source code is available at [github link will be added following acceptance].

